## Supplementary Materials and Methods for "Control of a gene transfer agent cluster in *Caulobacter crescentus* by transcriptional activation and anti-termination"

### Construction of plasmids and strains

***pNPTS138::gtaM-lacA***

The left flanking region (~500bp) of the insertion site, *lacA*, and the right flanking region (~500bp) of the insertion site were amplified by PCR using TLO3300+TLO3301, TLO3302+TLOO3303, TLO3304+TLOO3305 respectively. PCR products were purified from gel and assembled to BamHI-HindIII-cut pNPTS138 using 2x Gibson mastermix following the manufacturer's instruction. The Gibson reaction was used to transform chemically competent *E. coli* DH5α cells. The resulting plasmid was sequence verified by Sanger sequencing and used to transform *ΔlacA* competent cells to construct NTS2261.

***pNPTS138::flag-ihfA***

The left and right flanking regions (~500bp) of the insertion site were amplified by PCR using oligo NTO2830+NTO2831, NTO2832+NTO2833. PCR products were purified from gel and assembled to BamHI-HindIII-cut pNPTS138 using 2x Gibson mastermix following the manufacturer's instruction. The Gibson reaction was used to transform chemically competent *E. coli* DH5α cells. The resulting plasmid was sequence verified by Sanger sequencing and used to transform *C. crescentus* CB15N to make NTS2689.

***pNPTS138::ihfB-flag***

The left and right flanking regions (~500bp) of the insertion site were amplified by PCR using oligo using oligo NTO2745+NTO2746, NTO2747+NTO2748. PCR products were purified from gel and assembled to BamHI-HindIII-cut pNPTS138 using 2x Gibson mastermix following the manufacturer's instruction. The Gibson reaction was used to transform chemically competent *E. coli* DH5α cells. The resulting plasmid was sequence verified by Sanger sequencing and used to transform *C. crescentus* CB15N to make NTS2632.

***pNPTS138::ΔihfB***

The left and right flanking regions (~500bp) of the deletion site were amplified by PCR using oligo NTO2781+NTO2782, NTO2783+NTO2784. PCR products were purified from gel and assembled to BamHI-HindIII-cut pNPTS138 using 2x Gibson mastermix following the manufacturer's instruction. The Gibson reaction was used to transform chemically competent *E. coli* DH5α cells. The resulting plasmid was sequence verified by Sanger sequencing and used to transform *C. crescentus* CB15N to make NTS2608.

***pNPTS138::ΔihfA***

The left and right flanking regions (~500bp) of the deletion site were amplified by PCR using oligo NTO2799+NTO2800, NTO2801+NTO2802. PCR products were purified from gel and assembled to BamHI-HindIII-cut pNPTS138 using 2x Gibson mastermix following the manufacturer's instruction. The Gibson reaction was used to transform chemically competent *E. coli* DH5α cells. The resulting plasmid was sequence verified by Sanger sequencing and used to transform *C. crescentus* CB15N to make NTS2688.

***pXYFPC-1::ihfA***

*ihfA* orf was amplified by PCR from *C. crescentus* genomic DNA using NTO2845+NTO2846. PCR products were purified from gel and assembled to NdeI-NheI-cut pXYFPC-1 using 2x Gibson mastermix following the manufacturer's instructions. The Gibson reaction was used to transform chemically competent *E. coli* DH5α cells. The resulting plasmid was sequence verified by Sanger sequencing and used to transform NTS2692.

***pXYFPC-1::ihfB***
*ihfB* orf was amplified by PCR from *C. crescentus* genomic DNA using NTO2767+NTO2768. PCR products were purified from gel and assembled to NdeI-NheI-cut pXYFPC-1 using 2x Gibson mastermix following the manufacturer's instructions. The Gibson reaction was used to transform chemically competent *E. coli* DH5α cells. The resulting plasmid was sequence verified by Sanger sequencing and used to transform NTS2609.

***pNPTS138::ΔgtaT IBE***
To delete of 51bp spanning IHF binding element located in the promoter region of *gtaT* (*CCNA_02880*), the left and right flanking regions (~500bp) of the deletion site were amplified by PCR using oligo NTO2230+NTO2753, NTO2754+NTO2755. PCR products were purified from gel and assembled to BamHI-HindIII-cut pNPTS138 using 2x Gibson mastermix following the manufacturer's instruction. The Gibson reaction was used to transform chemically competent *E. coli* DH5α cells. The resulting plasmid was sequence verified by Sanger sequencing and used to transform *C. crescentus* CB15N to make NTS2623.

***pNPTS138::gtaT IBE (TT to GG)***
To change IHF binding element (IBE) in the promoter region of *gtaT* from TT to GG, the left and right flanking regions (~500bp) of the insertion site were amplified by PCR using oligo NTO2230+NTO2863, NTO2864+NTO2755. PCR products were purified from gel and assembled to BamHI-HindIII-cut pNPTS138 using 2x Gibson mastermix following the manufacturer's instruction. The Gibson reaction was used to transform chemically competent *E. coli* DH5α cells. The resulting plasmid was sequence verified by Sanger sequencing and used to transform NTS2623 to make NTS2724.

***pNPTS138::ΔgafY IBE***
To delete 42-bp spanning IHF binding element located in the promoter region of *gafY*, the left and right flanking regions (~500bp) of the deletion site were amplified by PCR using oligo NTO2749+NTO2750, NTO2751+NTO2752. PCR products were purified from gel and assembled to BamHI-HindIII-cut pNPTS138 using 2x Gibson mastermix following the manufacturer's instruction. The Gibson reaction was used to transform chemically competent *E. coli* DH5α cells. The resulting plasmid was sequence verified by Sanger sequencing and used to transform *C. crescentus* CB15N to make NTS2626.

***pNPTS138::gafY IBE (TT to GG)***
To change IHF binding element in the promoter region of *gafY* from TT to GG, the flanking regions of the insertion site were amplified by PCR using oligo 2749+2865, 2866+2752. PCR products were purified from gel and assembled to BamHI-HindIII-cut pNPTS138 using 2x Gibson mastermix following the manufacturer's instruction. The Gibson reaction was used to transform chemically competent *E. coli* DH5α cells. The resulting plasmid was sequence verified by Sanger sequencing and used to transform NTS2626 to make NTS2729.

***pNPTS138::Δihf500***
To delete 500bp spanning 4 IHF binding elements (IBE2-3-4-5) located inside the phage cluster, the left and right flanking regions (~500bp) of the deletion site were amplified by PCR using oligo NTO2888+NTO2889, NTO2890+NTO2891. PCR products were purified from gel and assembled to BamHI-HindIII-cut pNPTS138 using 2x Gibson mastermix following the manufacturer's instruction. The Gibson reaction was used to transform chemically competent *E. coli* DH5α cells. The resulting plasmid was sequence verified by Sanger sequencing and used to transform CB15N to make NTS2755.

***pNPTS138::ihfall4GG***
To change 4 IHF binding elements inside the phage cluster, the flanking regions of the insertion site were amplified by PCR using oligo NTO2888+NTO2892, NTO2893+NTO2929, NTO2932+NTO2935,

NTO2934+NTO2905, NTO2899+NTO2891. PCR products were purified from gel and assembled to BamHI-HindIII-cut pNPTS138 using 2x Gibson mastermix following the manufacturer's instruction. The Gibson reaction was used to transform chemically competent *E. coli* DH5α cells. The resulting plasmid was sequence verified by Plasmidsaurus and used to transform NTS2755 to make NTS2986.

***pNPTS138::nusG-vsvg***
To insert a VSVG tag at the C terminus of NusG, the flanking regions (~500bp) of the insertion site were amplified by PCR using oligo NTO2923+NTO2924, NTO2925+NTO2926. PCR products were purified from gel and assembled to BamHI-HindIII-cut pNPTS138 using 2x Gibson mastermix following the manufacturer's instruction. The Gibson reaction was used to transform chemically competent *E. coli* DH5α cells. The resulting plasmid was sequence verified by Sanger sequencing and used to transform CB15N to make NTS2758, and NTS2481 to make NTS2759, respectively.

***pNPTS138::vsvg-nusA***
To insert a VSVG tag at the N terminus of NusA, the flanking regions (~500bp) of the insertion site were amplified by PCR using oligo NTO4041+NTO4042, NTO4043+NTO4044. PCR products were purified from gel and assembled to BamHI-HindIII-cut pNPTS138 using 2x Gibson mastermix following the manufacturer's instruction. The Gibson reaction was used to transform chemically competent *E. coli* DH5α cells. The resulting plasmid was sequence verified by Sanger sequencing and used to transform CB15N to make NTS2897, and NTS2481 to make NTS2863, respectively.

***pNPTS138::vsvg-nusE***
To insert a VSVG tag at the N terminus of NusE, the flanking regions (~500bp) of the insertion site were amplified by PCR using oligo NTO4037+NTO4038, NTO4039+NTO4040. PCR products were purified from gel and assembled to BamHI-HindIII-cut pNPTS138 using 2x Gibson mastermix following the manufacturer's instruction. The Gibson reaction was used to transform chemically competent *E. coli* DH5α cells. The resulting plasmid was sequence verified by Sanger sequencing and used to transform CB15N to make NTS2901, and NTS2481 to make NTS2857, respectively.

***pNPTS138::flag-rpoD***
To insert a FLAG tag at the N terminus of RpoD, the flanking regions (~500bp) of the insertion site were amplified by PCR using oligo NTO4268+NTO4269, NTO4270+NTO4271. PCR products were purified from gel and assembled to BamHI-HindIII-cut pNPTS138 using 2x Gibson mastermix following the manufacturer's instruction. The Gibson reaction was used to transform chemically competent *E. coli* DH5α cells. The resulting plasmid was sequence verified by Sanger sequencing and used to transform CB15N to make NTS4195.

***pNTP2846***
To remove 2 BamHI sites from pXYFPC-1, using pXYFPC-1 as template, DNA fragments were amplified by PCR using NTO4188+NTO4191, NTO4189+NTO4190. PCR products were purified from gel and assembled using 2x Gibson mastermix following the manufacturer's instructions. The Gibson reaction was used to transform chemically competent *E. coli* DH5α cells. The resulting plasmid was sequence verified by Sanger sequencing.

***pNTP2864***
To remove the HindIII site from pNTP2846, using pNTPS2846 as template, DNA fragments were amplified by PCR using NTO4192+NTO4195, NTO4193+NTO4194. PCR products were purified from gel and assembled using 2x Gibson mastermix following the manufacturer's instructions. The Gibson reaction was used to transform chemically competent *E. coli* DH5α cells. The resulting plasmid was sequence verified by Sanger sequencing which is devoid of BamHI and HindIII sites.

***pRlacZ290:: PgtaT (-153 to +229)***

DNA fragment from –153 to +229 of GTA cluster promoter was amplified by PCR from genomic DNA using NTO4085+NTO4101. PCR products were purified from gel and assembled to EcoRI-KpnI-cut pRlacZ290 using 2x Gibson mastermix following the manufacturer's instruction. The Gibson reaction was used to transform chemically competent *E. coli* DH5α cells. The resulting plasmid was sequence verified by Sanger sequencing.

***pNTP2864:: PgtaT (-153 to +229)***

DNA fragment from –153 to +229 of GTA cluster promoter together with *lacZ* in pRlacZ290:: PgtaT (-153 to +229) was amplified by PCR using NTO4196+NTO4197. PCR products were purified from gel and assembled to NdeI-NheI-cut pNTP2864 using 2x Gibson mastermix following the manufacturer's instruction. The Gibson reaction was used to transform chemically competent *E. coli* DH5α cells. The resulting plasmid was sequence verified by whole-plasmid sequencing (Plasmidsaurus).

***pNPTS138:: PgtaT (-113 to +229)***

DNA fragment from –113 to +229 of GTA cluster promoter was amplified by PCR from genomic DNA using NTO4259+NTO4185. PCR products were purified from gel and assembled to BamHI-HindIII-cut pNPTS138 using 2x Gibson mastermix following the manufacturer's instruction. The Gibson reaction was used to transform chemically competent *E. coli* DH5α cells. The resulting plasmid was sequence verified by Sanger sequencing.

***pNTP2864:: PgtaT (-113 to +229)***

DNA fragment from –113 to +229 of GTA cluster promoter was removed from pNPTS138:: PgtaT (-113 to +229) and ligated to EcoRI-KpnI-cut pNTP2864:: PgtaT (-153 to +229). The ligation mixture was used to transform chemically competent *E. coli* DH5α cells. The resulting plasmid was verified by restriction digestion.

***pNPTS138:: PgtaT (-56 to +229)***

DNA fragment from –56 to +229 of GTA cluster promoter was amplified by PCR from genomic DNA using NTO4369+NTO4185. PCR products were purified from gel and assembled to BamHI-HindIII-cut pNPTS138 using 2x Gibson mastermix following the manufacturer's instruction. The Gibson reaction was used to transform chemically competent *E. coli* DH5α cells. The resulting plasmid was sequence verified by Sanger sequencing.

***pNTP2864:: PgtaT (-56 to +229)***

DNA fragment from –56 to +229 of GTA cluster promoter was removed from pNPTS138:: PgtaT (-56 to +229) and ligated to EcoRI-KpnI-cut pNTP2864:: PgtaT (-153 to +229). The ligation mixture was used to transform chemically competent *E. coli* DH5α cells. The resulting plasmid was verified by restriction digestion.

***pNPTS138:: PgtaT (-143 to +229)***

DNA fragment from –143 to +229 of GTA cluster promoter was amplified by PCR from genomic DNA using NTO4256+NTO4185. PCR products were purified from gel and assembled to BamHI-HindIII-cut pNPTS138 using 2x Gibson mastermix following the manufacturer's instruction. The Gibson reaction was used to transform chemically competent *E. coli* DH5α cells. The resulting plasmid was sequence verified by Sanger sequencing.

***pNTP2864:: PgtaT (-143 to +229)***

DNA fragment from –143 to +229 of GTA cluster promoter was removed from pNPTS138:: PgtaT (-143 to +229) and ligated to EcoRI-KpnI-cut pNTP2864:: PgtaT (-153 to +229). The ligation mixture was used to transform chemically competent *E. coli* DH5α cells. The resulting plasmid was verified by restriction digestion.

***pNPTS138:: PgtaT (-133 to +229)***

DNA fragment from –133 to +229 of GTA cluster promoter was amplified by PCR from genomic DNA using NTO4257+NTO4185. PCR products were purified from gel and assembled to BamHI-HindIII-cut pNPTS138 using 2x Gibson mastermix following the manufacturer's instruction. The Gibson reaction was used to transform chemically competent *E. coli* DH5α cells. The resulting plasmid was sequence verified by Sanger sequencing.

***pNTP2864:: PgtaT (-133 to +229)***

DNA fragment from –133 to +229 of GTA cluster promoter was removed from pNPTS138:: PgtaT (-133 to +229) and ligated to EcoRI-KpnI-cut pNTP2864:: PgtaT (-153 to +229). The ligation mixture was used to transform chemically competent *E. coli* DH5α cells. The resulting plasmid was verified by restriction digestion.

***pNPTS138:: PgtaT (-123 to +229)***

DNA fragment from –123 to +229 of GTA cluster promoter was amplified by PCR from genomic DNA using NTO4258+NTO4185. PCR products were purified from gel and assembled to BamHI-HindIII-cut pNPTS138 using 2x Gibson mastermix following the manufacturer's instruction. The Gibson reaction was used to transform chemically competent *E. coli* DH5α cells. The resulting plasmid was sequence verified by Sanger sequencing.

***pNTP2864::PgtaT (-123 to +229)***

DNA fragment from –123 to +229 of GTA cluster promoter was removed from pNPTS138:: PgtaT (-123 to +229) and ligated to EcoRI-KpnI-cut pNTP2864:: PgtaT (-153 to +229). The ligation mixture was used to transform chemically competent *E. coli* DH5α cells. The resulting plasmid was verified by restriction digestion.

***pNTPS138:: Δ125bp (+90 to +214)***

To delete 125bp in gtaT to find the correct start codon, the flanking regions (~500bp) of the deletion site were amplified by PCR using NTO2230+NTO2764, NTO2765+NTO2766. PCR products were purified from gel and assembled to BamHI-HindIII-cut pNPTS138 using 2x Gibson mastermix following the manufacturer's instruction. The Gibson reaction was used to transform chemically competent *E. coli* DH5α cells. The resulting plasmid was sequence verified by Sanger sequencing, and introduced into CB15N to make NTS2653.

***pNTPS138:: Δ144bp (+230 to 373)***

To delete 144bp in *gtaT* to find the correct start codon, the flanking regions (~500bp) of the deletion site were amplified by PCR using NTO2957+NTO2958, NTO2960+NTO2961. PCR products were purified from gel and assembled to BamHI-HindIII-cut pNPTS138 using 2x Gibson mastermix following the manufacturer's instruction. The Gibson reaction was used to transform chemically competent *E. coli* DH5α cells. The resulting plasmid was sequence verified by Sanger sequencing, and introduced into CB15N to make NTS2792.

***pNPTS138::gtaT (ATG1 to TGA)***

To mutagenize the $1^{st}$ ATG codon to TGA, the flanking regions (~500bp) of the insertion site were amplified by PCR using NTO2957+NTO2962, NTO2963+NTO2961. PCR products were purified from gel and assembled to BamHI-HindIII-cut pNPTS138 using 2x Gibson mastermix following the manufacturer's instruction. The Gibson reaction was used to transform chemically competent *E. coli* DH5α cells. The resulting plasmid was sequence verified by Sanger sequencing, and introduced into NTS2792 to make NTS2800.

***pNPTS138:: gtaT (ATG2 to TGA)***

To mutagenize the 1st ATG codon to TGA, the flanking regions (~500bp) of the insertion site were amplified by PCR using NTO2957+NTO2964, NTO2965+NTO2961. PCR products were purified from gel and assembled to BamHI-HindIII-cut pNPTS138 using 2x Gibson mastermix following the manufacturer's instruction. The Gibson reaction was used to transform chemically competent *E. coli* DH5α cells. The resulting plasmid was sequence verified by Sanger sequencing, and introduced into NTS2792 to make NTS2801.

***pNPTS138:: gtaT (ATG3 to TGA)***

To mutagenize the 1st ATG codon to TGA, the flanking regions (~500bp) of the insertion site were amplified by PCR using NTO2957+NTO2966, NTO2967+NTO2961. PCR products were purified from gel and assembled to BamHI-HindIII-cut pNPTS138 using 2x Gibson mastermix following the manufacturer's instruction. The Gibson reaction was used to transform chemically competent *E. coli* DH5α cells. The resulting plasmid was sequence verified by Sanger sequencing, and introduced into NTS2792 to make NTS2802.

***pNPTS138::Δ336bp (-113 to +223)***

To delete 336bp from the putative start codon TTG of gtaT to the putative RBS, the flanking regions (~500bp) of the deletion site were amplified by PCR using NTO2230+NTO2906, NTO2907+NTO2766. PCR products were purified from gel and assembled to BamHI-HindIII-cut pNPTS138 using 2x Gibson mastermix following the manufacturer's instruction. The Gibson reaction was used to transform chemically competent *E. coli* DH5α cells. The resulting plasmid was sequence verified by Sanger sequencing, and introduced into CB15N to make NTS2757.

***pNPTS138::gtaT (TTG to CTG)***

To change the annotated start codon of gtaT from TTG to CTG, the flanking regions (~500bp) of the insertion site were amplified by PCR using NTO2230+NTO2908, NTO2909+NTO2766. PCR products were purified from gel and assembled to BamHI-HindIII-cut pNPTS138 using 2x Gibson mastermix following the manufacturer's instruction. The Gibson reaction was used to transform chemically competent *E. coli* DH5α cells. The resulting plasmid was sequence verified by Sanger sequencing, and introduced into NTS2757 to make NTS2767.

***pNPTS138::ΔYBE***

To delete GafY binding element (YBE) in GTA cluster promoter, the flanking regions (~500bp) of the deletion site were amplified by PCR using NTO2230+NTO2970, NTO2971+NTO2755. PCR products were purified from gel and assembled to BamHI-HindIII-cut pNPTS138 using 2x Gibson mastermix following the manufacturer's instruction. The Gibson reaction was used to transform chemically competent *E. coli* DH5α cells. The resulting plasmid was sequence verified by Sanger sequencing, and introduced into CB15N to make NTS2799.

***pNPTS138:: YBE* (ACTATG to TGACCC)***

To change GafY binding element in GTA cluster promoter from ACTATGGGGA to tgacccGGGA, the flanking regions (~500bp) of the insertion site were amplified by PCR using NTO2230+NTO4157, NTO4158+NTO2755. PCR products were purified from gel and assembled to BamHI-HindIII-cut pNPTS138 using 2x Gibson mastermix following the manufacturer's instruction. The Gibson reaction was used to transform chemically competent *E. coli* DH5α cells. The resulting plasmid was sequence verified by Sanger sequencing, and introduced into NTS2799 to make NTS2940.

***pNPTS138::ΔGTA core promoter***

To delete the core promoter of GTA cluster, the flanking regions (~500bp) of the deletion site were amplified by PCR using NTO2230+NTO2760, NTO2761+NTO2755. PCR products were purified from gel and assembled to BamHI-HindIII-cut pNPTS138 using 2x Gibson mastermix following the manufacturer's instruction. The Gibson reaction was used to transform chemically competent *E. coli* DH5α cells. The resulting plasmid was sequence verified by Sanger sequencing, and introduced into CB15N to make NTS2650.

***pNPTS138::ZBE* (CTGGCGC to TTTTTTC)***

To change GafZ binding element (ZBE) from GCGCCCGCTGGCGC to GCGCCCGTTTTTTC, the flanking regions (~500bp) of the insertion site were amplified by PCR using NTO2230+NTO4090, NTO4091+NTO2755. PCR products were purified from gel and assembled to BamHI-HindIII-cut pNPTS138 using 2x Gibson mastermix following the manufacturer's instruction. The Gibson reaction was used to transform chemically competent *E. coli* DH5α cells. The resulting plasmid was sequence verified by Sanger sequencing, and introduced into NTS2650 to make NTS2955.

***pNPTS138::mut10-1***

To change the -10 promoter sequence of GTA cluster promoter from TATCTTT to ccccccc, PCR amplifies fragments using NTO4184+NTO4174, NTO4175+NTO4185. PCR products were purified from gel and assembled to BamHI-HindIII-cut pNPTS138 using 2x Gibson mastermix following the manufacturer's instruction. The Gibson reaction was used to transform chemically competent *E. coli* DH5α cells. The resulting plasmid was sequence verified by Sanger sequencing.

***pNTP2864::mut10-1***

GTA cluster promoter with the -10 sequence changed from TATCTTT to ccccccc was removed from pNPTS138::mut10-1 as EcoRI-KpnI fragment and ligated to EcoRI-KpnI-cut pNTP2864:: PgtaT (-153 to +229) The ligation mixture was used to transform chemically competent *E. coli* DH5α cells. The resulting plasmid was verified by restriction digestion.

***pNPTS138::mut10-2***

To change the -10 promoter sequence of GTA cluster promoter from TATCTTTGT to TATCTcccc, PCR amplifies fragments using NTO4184+NTO4176, NTO4177+NTO4185. PCR products were purified from gel and assembled to BamHI-HindIII-cut pNPTS138 using 2x Gibson mastermix following the manufacturer's instruction. The Gibson reaction was used to transform chemically competent *E. coli* DH5α cells. The resulting plasmid was sequence verified by Sanger sequencing.

***pNTP2864::mut10-2***

GTA cluster promoter with -10 sequence changed from TATCTTTGT to TATCTcccc was removed from pNPTS138::mut10-2 as EcoRI-KpnI fragment and ligated to EcoRI-KpnI-cut pNTP2864:: PgtaT (-153 to +229). The ligation mixture was used to transform chemically competent *E. coli* DH5α cells. The resulting plasmid was verified by restriction digestion.

***pNPTS138::mut35-1***

To change the -35 promoter sequence of GTA cluster promoter from AATCAT to cccccc, PCR amplifies fragments using NTO4184+NTO4242, NTO4243+NTO4185. PCR products were purified from gel and assembled to BamHI-HindIII-cut pNPTS138 using 2x Gibson mastermix following the manufacturer's instruction. The Gibson reaction was used to transform chemically competent *E. coli* DH5α cells. The resulting plasmid was sequence verified by Sanger sequencing.

***pNTP2864::mut35-1***

GTA cluster promoter with the -35 sequence changed from AATCAT to cccccc was removed from pNPTS138::mut35-1 as EcoRI-KpnI fragment and ligated to EcoRI-KpnI-cut pNTP2864:: PgtaT (-153 to +229). The ligation mixture was used to transform chemically competent *E. coli* DH5α cells. The resulting plasmid was sequence verified by Sanger sequencing.

***pNPTS138::mut35-2***
To change the -35 promoter sequence of GTA cluster promoter from AATCAT to AAgggT, PCR amplifies fragments using NTO4184+NTO4264, NTO4265+NTO4185. PCR products were purified from gel and assembled to BamHI-HindIII-cut pNPTS138 using 2x Gibson mastermix following the manufacturer's instruction. The Gibson reaction was used to transform chemically competent *E. coli* DH5α cells. The resulting plasmid was sequence verified by Sanger sequencing.

***pNTP2864::mut35-2***
GTA cluster promoter with the -35 sequence changed from AATCAT to AAgggT was removed from pNPTS138::mut35-2 as EcoRI-KpnI fragment and ligated to EcoRI-KpnI-cut pNTP2864:: PgtaT (-153 to +229). The ligation mixture was used to transform chemically competent *E. coli* DH5α cells. The resulting plasmid was sequence verified by Sanger sequencing.

***pNPTS138::PrsaA***
PrsaA was amplified by PCR from *C. crescentus* genomic DNA using NTO4252+NTO4253. PCR products were purified from gel and assembled to BamHI-HindIII-cut pNPTS138 using 2x Gibson mastermix following the manufacturer's instruction. The Gibson reaction was used to transform chemically competent *E. coli* DH5α cells. The resulting plasmid was sequence verified by Sanger sequencing.

***pNTP2864:: PrsaA***
PrsaA was removed from pNPTS138:: PrsaA as EcoRI-KpnI fragment and ligated to EcoRI-KpnI-cut pNTP2864:: PgtaT (-153 to +229). The ligation mixture was used to transform chemically competent *E. coli* DH5α cells. The resulting plasmid was verified by restriction digestion.

***pNPTS138::terGTA***
Using genomic DNA as template, DNA fragments were amplified by PCR using NTO4094+NTO2930, NTO2931+NTO2935, NTO2936+NTO4095. PCR products were purified from gel and assembled to BamHI-HindIII-cut pNPTS138 using 2x Gibson mastermix following the manufacturer's instruction. The Gibson reaction was used to transform chemically competent *E. coli* DH5α cells. The resulting plasmid was sequence verified by whole-plasmid sequencing (Plasmidsaurus).

***pNTP2884:: terGTA***
terGTA was removed from pNPTS138::terGTA as BamHI-HindIII fragment and ligated to BamHI-HindIII cut pNTP2884. The ligation mixture was used to transform chemically competent *E. coli* DH5α cells. The resulting plasmid was verified by restriction digestion.

***pNTP2864:: PgtaT (-153 to +229) + terGTA***
terGTA was removed from pNPTS138::terGTA as a BamHI-HindIII fragment and ligated to BamHI-HindIII cut pNTP2864:: PgtaT (-153 to +229). The ligation mixture was used to transform chemically competent *E. coli* DH5α cells. The resulting plasmid was verified by whole-plasmid sequencing (Plasmidsaurus).

***pNTP2864::PrsaA + terGTA***
PrsaA promoter was removed from pNPTS138:: PrsaA as an EcoRI-KpnI fragment and ligated to EcoRI-KpnI-cut pNTP2864:: PgtaT (-153 to +229) + terGTA. The ligation mixture was used to transform chemically competent *E. coli* DH5α cells. The resulting plasmid was verified by Plasmidsaurus.

***pNTP2884***

pNTP2864:: PgtaT (-153 to +229) was cut with EcoRI and KpnI to remove the PgtaT-153 to +229, blunted with DNA polymerase I (NEB) and re-ligated using T4 ligase (NEB). The ligation mixture was used to transform chemically competent *E. coli* DH5α cells. The resulting plasmid was verified by Sanger sequencing.

***pCOLA-Duet1::6xhis gafZ***
*gafZ* was amplified by PCR from genomic DNA using NTO2693+NTO2694. PCR products were purified from gel and assembled to EcoRI-HindIII cut pCOLA-Duet1 using 2x Gibson mastermix following the manufacturer's instruction. The Gibson reaction was used to transform chemically competent *E. coli* DH5α cells. The resulting plasmid was sequence verified by Sanger sequencing.

***pCOLA-Duet1::6xhis gafZ gafY***
*gafY* was amplified by PCR from genomic DNA using NTO2691+NTO2692. PCR products were purified from gel and assembled to NdeI-KpnI cut pCOLA-Duet1::6xhis gafZ using 2x Gibson mastermix following the manufacturer's instruction. The Gibson reaction was used to transform chemically competent *E. coli* DH5α cells. The resulting plasmid was sequence verified by Sanger sequencing.

***pCOLADuet-1::6xhis rpoD gafY***
*rpoD* was amplified by PCR from genomic DNA using NTO4289+NTO4290. PCR products were purified from gel and assembled to EcoRI-HindIII cut pCOLA-Duet1::6xhis gafZ gafY using 2x Gibson mastermix following the manufacturer's instruction. The Gibson reaction was used to transform chemically competent *E. coli* DH5α cells. The resulting plasmid was sequence verified by Sanger sequencing.

***pCOLADuet-1::6xhis rpoD (domain 3+4)***
pCOLADuet-1::6xhis rpoD gafY was cut with NdeI and KpnI to remove the *gafY* insert. The backbone containing 6xhis rpoD was treated with Klenow to blunt the ends and ligated using T4 ligase. The ligation mixture was used to transform chemically competent *E. coli* DH5α cells. The resulting plasmid was purified and verified by Sanger sequencing. The plasmid was subsequently cut with EcoRI-HindIII to remove the *rpoD* insert. The newly generated backbone was ligated to EcoRI-HindIII cut rpoD3+4 fragment which was amplified by PCR using NTO4349+NTO4290. The ligation reaction was used to transform chemically competent *E. coli* DH5α cells. The resulting plasmid was sequence verified by Sanger sequencing.

***pET15b::gafY***
*gafY* orf was amplified by PCR using NTO4346 and NTO4347. PCR products were purified from gel and assembled to NcoI-BamHI-cut pET15b using 2x Gibson mastermix following the manufacturer's instruction. The Gibson reaction was used to transform chemically competent *E. coli* DH5α cells. The resulting plasmid was sequence verified by Sanger sequencing.

***pNPTS138:: ΔgafZ***
To delete *gafZ*, the flanking regions (~500bp) of the deletion site were amplified by PCR using NTO2634+NTO2635, NTO2636+NTO2637. PCR products were purified from gel and assembled to BamHI-HindIII-cut pNPTS138 using 2x Gibson mastermix following the manufacturer's instruction. The Gibson reaction was used to transform chemically competent *E. coli* DH5α cells. The resulting plasmid was sequence verified by Sanger sequencing, and introduced into *ΔlacA* cells to make NTS3086.

***pNPTS138::ΔrogA***
To delete *rogA*, the flanking regions (~500bp) of the deletion site were amplified by PCR using NTO2194+NTO2195, NTO2196+NTO2197. PCR products were purified from gel and assembled to BamHI-HindIII-cut pNPTS138 using 2x Gibson mastermix following the manufacturer's instruction. The Gibson reaction was used to transform chemically competent *E. coli* DH5α cells. The resulting plasmid

was sequence verified by Sanger sequencing, and introduced into *ΔlacA* cells to make NTS3011, into NTS3086 to make NTS3124.

***pNPTS138::terT1***

terT1 was amplified by PCR from pBXMCS-2 using NTO4151 and NTO4152. PCR products were purified from gel and assembled to BamHI-HindIII-cut pNPTS138 using 2x Gibson mastermix following the manufacturer's instruction. The Gibson reaction was used to transform chemically competent *E. coli* DH5α cells. The resulting plasmid was sequence verified by Sanger sequencing.

***pNTP2864:: PgtaT (-153 to +229) + terT1***

terT1 was restricted out from pNPTS138::terT1 as a BamHI-HindIII-cut fragment and ligated to BamHI-HindIII-cut pNTP2864::PgtaT (-153 to +229). The ligation reaction was used to transform chemically competent *E. coli* DH5α cells. The resulting plasmid was sequence verified by Sanger sequencing.

***pNTP2884:: terT1***

pNTP2864:: PgtaT (-153 to +229) + terT1 was cut with EcoRI-KpnI to remove the PgtaT (-153 to +229), blunted with DNA polymerase I (NEB) and re-ligated using T4 ligase (NEB). The ligation mixture was used to transform chemically competent *E. coli* DH5α cells. The resulting plasmid was verified by Sanger sequencing.

***pNTP2864::PrsaA + terT1***

PrsaA was removed from pNPTS138::PrsaA as an EcoRI-KpnI fragment and ligated to EcoRI-KpnI-cut pNTP2864::PgtaT (-153 to +229) + terT1. The ligation reaction was used to transform chemically competent *E. coli* DH5α cells. The resulting plasmid was sequence verified by Sanger sequencing.

**Supplementary Table 1.** CDS and their annotated functions on the main CcGTA cluster

| start nt | end nt | strand | length (amino acids) | CDS | locus tag | conventional name | Annotations |
|---|---|---|---|---|---|---|---|
| 3013149 | 3015680 | - | 843 | CDS | CCNA_02861 | *gtaB* | phage host specificity protein |
| 3015316 | 3016860 | - | 514 | CDS | CCNA_02862 | *gatC* | phage host specificity protein |
| 3016985 | 3017605 | - | 206 | CDS | CCNA_02863 | *gtaD* | Spr-family cell wall-associated hydrolase |
| 3017602 | 3018228 | - | 208 | CDS | CCNA_02864 | *gtaE* | phage conserved hypothetical protein |
| 3018228 | 3018863 | - | 211 | CDS | CCNA_02865 | *gtaF* | phage-derived conserved hypothetical protein |
| 3019011 | 3019475 | + | 154 | CDS | CCNA_02866 | - | hypothetical protein |
| 3019649 | 3020155 | - | 168 | CDS | CCNA_02867 | *gtaG* | phage tail length tape measure-related protein |
| 3020152 | 3020331 | - | 59 | CDS | CCNA_02868 | *gtaH* | hypothetical protein |
| 3020328 | 3020612 | - | 94 | CDS | CCNA_02869 | *gtaI* | hypothetical protein |
| 3020768 | 3021184 | - | 138 | CDS | CCNA_02870 | *gtaJ* | phage protein |
| 3021200 | 3021598 | - | 132 | CDS | CCNA_02871 | *gtaK* | hypothetical protein |
| 3021595 | 3021876 | - | 93 | CDS | CCNA_03882 | *gtaL* | phage gp6-like head-tail connector protein |
| 3022019 | 3023239 | - | 406 | CDS | CCNA_02872 | *gtaM* | phage major capsid |

| | | | | | | | |
|---|---|---|---|---|---|---|---|
| 3023436 | 3023798 | + | 120 | CDS | CCNA_02873 | - | glyoxalase/bleomycin resistance protein, dioxygenase superfamily |
| 3023772 | 3024188 | - | 138 | CDS | CCNA_02874 | *gtaN* | phage prohead protease |
| 3024178 | 3024576 | - | 132 | CDS | CCNA_02875 | *gtaO* | hypothetical protein |
| 3024589 | 3025917 | - | 442 | CDS | CCNA_02877 | *gtaP* | portal protein |
| 3025889 | 3026329 | + | 146 | CDS | CCNA_02876 | | very short patch repair (Vsr) endonuclease |
| 3026557 | 3026715 | - | 52 | CDS | CCNA_02878 | *gtaQ* | hypothetical protein |
| 3026790 | 3026885 | - | 31 | CDS | CCNA_02879 | *gtaS* | genetic exchange related protein |
| 3027016 | 3028719 | - | 567 | CDS | CCNA_02880 | *gtaT* | terminase-like family protein |

**Supplementary Table 2.** Strains, plasmids, and oligonucleotides

| **Strains** | **Strains/Descriptions** | **Sources** |
|---|---|---|
| Rosetta (DE3) | *E. coli* host for protein overexpression from an IPTG-inducible T7 promoter | Merck |
| CB15N | NA1000, wild-type *C. crescentus* | Lab collection |
| NTS2975 | Δ*lacA* | [1] |
| NTS2261 | Δ*lacA CCNA_02872 (gtaM)::gtaM-lacA* | This study |
| NTS2275 | CB15N Δ*rogA::tetracyclineR* | This study |
| NTS2305 | CB15N Δ*rogA::tetracyclineR* Δ*lacA gtaM::gtaM-lacA* | This study |
| NTS2481 | CB15N *gafZ::flag-gafZ* | This study |
| NTS2501 | CB15N Δ*gafY* | This study |
| NTS2515 | CB15N Δ*rogA::tetracyclineR* Δ*gafY* | This study |
| NTS2489 | CB15N Δ*rogA::tetracyclineR gafZ::flag-gafZ* | This study |
| NTS2689 | CB15N *ihfA::flag-ihfA* | This study |
| NTS2696 | CB15N Δ*rogA::tetracyclineR ihfA::flag- ihfA* | This study |
| NTS2632 | CB15N *ihfB:: ihfB-flag* | This study |
| NTS2638 | CB15N Δ*rogA::tetracyclineR ihfB:: ihfB-flag* | This study |
| NTS2688 | CB15N Δ*ihfA* | This study |
| NTS2692 | CB15N Δ*rogA::tetracyclineR* Δ*ihfA* | This study |
| NTS2608 | CB15N Δ*ihfB* | This study |
| NTS2609 | CB15N Δ*rogA::tetracyclineR* Δ*ihfB* | This study |
| NTS2623 | CB15N Δ*gtaT IBE* | This study |
| NTS2724 | CB15N *gtaT IBE1 (TT to GG)* | This study |
| NTS2730 | CB15N Δ*rogA::tetracyclineR gtaT IBE1 (TT to GG)* | This study |
| NTS2626 | CB15N Δ*gafY IBE6* | This study |
| NTS2729 | CB15N *gafY IBE6 (TT to GG)* | This study |

| | | |
|---|---|---|
| NTS2735 | CB15N Δ*rogA::tetracyclineR gafY IBE6 (TT to GG)* | This study |
| NTS2755 | CB15N Δ*ihf500* | This study |
| NTS2986 | CB15N ihfall4GG | This study |
| NTS3038 | CB15N Δ*rogA::tetracyclineR* ihfall4GG | This study |
| | CB15N β' RNAP:: 3xflag-β' RNAP | Gift from M. Laub |
| NTS2640 | CB15N Δ*rogA::tetracyclineR* β' RNAP:: flag- β' RNAP | This study |
| NTS2758 | CB15N *nusG::nusG-VSVG* | This study |
| NTS2781 | CB15N Δ*rogA::tetracyclineR nusG::nusG-VSVG* | This study |
| NTS2759 | CB15N *nusG::nusG-VSVG gafZ::flag-gafZ* | This study |
| NTS2782 | CB15N *ΔrogA::tetracyclineR nusG::nusG-VSVG gafZ::flag-gafZ* | This study |
| NTS2897 | CB15N *nusA::VSVG-nusA* | This study |
| NTS2903 | CB15N *ΔrogA::tetracyclineR nusA::VSVG-nusA* | This study |
| NTS2863 | CB15N *nusA::VSVG-nusA gafZ::flag-gafZ* | This study |
| NTS2869 | CB15N *ΔrogA::*tetracyclineR nusA::VSVG-nusA *gafZ::flag-gafZ* | This study |
| NTS2901 | CB15N *nusE::VSVG-nusE* | This study |
| NTS2907 | CB15N Δ*rogA::tetracyclineR nusE::VSVG-nusE* | This study |
| NTS2857 | CB15N *nusE::VSVG-nusE gafZ::flag-gafZ* | This study |
| NTS2871 | CB15N *ΔrogA::tetracyclineR nusE::VSVG-nusE gafZ::flag-gafZ* | This study |
| NTS4195 | CB15N *rpoD::flag-rpoD* | This study |
| NTS2519 | CB15N Δ*rogA::tetracyclineR rpoD::flag-rpoD* | This study |
| NTS2650 | CB15NΔ*GTAcore promoter* | This study |
| NTS2955 | CB15N *ZBE** (CTGGCGC to TTTTTTC)) | This study |
| NTS2970 | CB15N *ΔrogA::tetracyclineR ZBE** (CTGGCGC to TTTTTTC) | This study |
| NTS2799 | CB15N Δ*YBE at the GTA promoter* | This study |
| NTS2940 | CB15N *YBE* (ACTATG* to *TGACCC)* | This study |
| NTS3169 | CB15N *ΔrogA::tetracyclineR YBE* (ACTATG* to *TGACCC)* | This study |
| NTS3011 | CB15N Δ*lacA* Δ*rogA* | This study |
| NTS3086 | CB15N Δ*lacA* Δ*gafZ* | This study |
| NTS3124 | CB15N Δ*lacA* Δ*rogA* Δ*gafZ* | This study |
| NTS2653 | CB15N *Δ125bp* (+90 to +214) | This study |
| NTS2757 | CB15N *Δ336bp (-113 to +223)* | This study |
| NTS2792 | CB15N Δ*144bp (+230 to 373)* | This study |
| NTS2767 | CB15N *gtaT TTG to CTG* (using NTS2757 as background) | This study |
| NTS2777 | CB15N Δ*rogA::tetracyclineR gtaT (TTG to CTG)* | This study |
| NTS2800 | CB15N *gtaT ATG1 to TGA* (using NTS2792 as background) | This study |

| | | |
|---|---|---|
| NTS2811 | CB15N Δ*rogA::tetracyclineR gtaT (ATG1 to TGA)* | This study |
| NTS2801 | CB15N *gtaT ATG2 to TGA* (using NTS2792 as background) | This study |
| NTS2812 | CB15N Δ*rogA::tetracyclineR gtaT (ATG2 to TGA)* | This study |
| NTS2802 | CB15N *gtaT ATG3 to TGA* (using NTS2792 as background) | This study |
| NTS2809 | CB15N Δ*rogA::tetracyclineR gtaT (ATG3 to TGA)* | This study |

| **Plasmids** | **Descriptions** | **Source** |
|---|---|---|
| pNPTS138 | suicide vector for gene knock-in/knock-out in *C. crescentus*, *sacB*, kanamycinR | Lab collection |
| pNPTS138::*CCNA_02872-lacA* | For transcriptional fusion of *lacA* to the C-terminus of *gtaM* | This study |
| pNPTS138::*flag-ihfA* | For the clean insertion of FLAG tag to the N-terminus of *ihfA* | This study |
| pNPTS138::*ihfB-flag* | For the clean insertion of FLAG tag to the C-terminus of *ihfB* | This study |
| pNPTS138::Δ*ihfA* | For the deletion of *ihfA* ORF | This study |
| pNPTS138::Δ*ihfB* | For the deletion of *ihfB* ORF | This study |
| pXYFPC-1 | Integrative vector, for insertion at the *xyl* locus, spectinomycinR | [2] |
| pBXMCS-2 | To amplify $ter_{T1}$ | 2 |
| pXYFPC-1:: *ihfA* | For complementation of Δ*ihfA*, ectopic expression of *ihfA* from the *xyl* locus, xylose inducible | This study |
| pXYFPC-1:: *ihfB* | For complementation of Δ*ihfB,* ectopic expression of *ihfB* from the *xyl* locus, xylose inducible | This study |
| ***pNPTS138::ΔgtaT IBE*** | For the deletion of 51 bp spanning IHF binding element located in the promoter region of *gtaT* | This study |
| ***pNPTS138::gtaT IBE*** *(TT to GG)* | For the mutagenesis of IHF binding element in the promoter region of *gtaT* from TT to GG | This study |
| pNPTS138::Δ*gafY IBE* | For the deletion of 42 bp spanning IHF binding element located in the promoter region of *gafY* | This study |
| pNPTS138::*gafY IBE (TT to GG)* | For the mutagenesis of IHF binding element in the promoter region of *gafY* from TT to GG | This study |
| pNPTS138::Δ*ihf500* | For the deletion of 500 bp spanning 4 IHF binding sites (IBE2-3-4-5) located inside the phage cluster | This study |
| pNPTS138::*ihfall4GG* | For the mutagenesis of all 4 IHF binding elements located inside the phage cluster | This study |
| pRlacZ290 | Transcriptional *lacZ* reporter plasmid, tetracyclineR | Gift from Clare Kirkpatrick |

| | | |
|---|---|---|
| pNTP2846 | Same as pXYFPC-1 but the BamHI site was removed | This study |
| pNTP2864 | Same as pNTP2846 but the HindIII site was removed | This study |
| pNPTS138::*nusG-vsvg* | For a clean insertion of VSVG tag to the C-terminus of NusG | This study |
| pNPTS138::*vsvg-nusA* | For a clean insertion of VSVG tag to the N-terminus of NusA | This study |
| pNPTS138::*vsvg-nusE* | For a clean insertion of VSVG tag to the N-terminus of NusE | This study |
| pNPTS138::*flag-rpoD* | For a clean insertion of a FLAG tag to the N-terminus of RpoD | This study |
| ***pNPTS138::ΔYBE*** | For deletion of YBE in *gtaT* promoter | This study |
| ***pNPTS138::YBE* (ACTATGto TGACCC)*** | For mutagenesis of YBE from ACTATG to TGACCC | This study |
| ***pNPTS138::ΔGTAcore promoter*** | For deletion of the core promoter of *gtaT* CGCTGGCGCCGTATCTTTGTGTCATGGCCG GAAGAAGCG | This study |
| ***pNPTS138::ZBE* (CTGGCGC to TTTTTTC)*** | For mutagenesis of ZBE from CTGGCG to TTTTTT | This study |
| pNPTS138::*Δ125bp* (+90 to +214) | To identify the correct ORF of *gtaT* | This study |
| pNPTS138::*Δ336bp (-113 to +223)* | To change putative start codon of *gtaT* from TTG to CTG | This study |
| pNPTS138::*gtaT TTG to CTG* | To change putative start codon of *gtaT* from TTG to CTG | |
| pNPTS138::*Δ144bp(+230 to 373)* | To change putative start codon of *gtaT* from ATGs to TGA | This study |
| *pNPTS138::gtaT (ATG1 to TGA)* | To change the putative start codon ATG1 to TGA | This study |
| pNPTS138::*gtaT (ATG2 to TGA)* | To change the putative start codon ATG2 to TGA | This study |
| pNPTS138::*gtaT (ATG3 to TGA)* | To change the putative start codon ATG3 to TGA | This study |
| pCOLA-Duet1 | For protein overexpression from an IPTG inducible T7 promoter, kanamycinR | Merck |
| pET15b | For protein overexpression from an IPTG inducible T7 promoter, carbenicillinR | Merck |
| pCOLA-Duet1::6x*his gafZ* | Overexpression of 6xHis GafZ (cloned into the MCS-1) of pCOLA-Duet1 | This study |
| pCOLA-Duet1::6x*his gafZ gafY* | Overexpression of 6xHis GafZ (cloned into the MCS-1) and GafY (cloned into the MCS-2) in pCOLA-Duet1 | This study |

| | | |
|---|---|---|
| pCOLA-Duet1:: *6xhis rpoD gafY* | Overexpression 6xHis RpoD (cloned into the MCS-1) and GafY (cloned into the MCS-2) in pCOLA-Duet1 | This study |
| pCOLA-Duet1:: 6x*his rpoD3+4* | Overexpression of 6xHis RpoD domain 3+4 (cloned into the MCS-1) in pCOLA-Duet1 | This study |
| pET15b::*gafY* | Overexpression of GafY in pET15b | This study |
| pRlacZ290:: PgtaT-*153 to +229* | Sequence from -153 to +229 of the GTA cluster promoter PgtaT cloned in pRlacZ290 | This study |
| pNTP2864:: PgtaT (-153 to +229) | Sequence from -153 to +229 of the GTA cluster promoter fused to *lacZ* was amplified from pRlacZ290::-153 to +229 PgtaT, and was subsequently cloned in between the NdeI and NheI sites of pNTP2864 | This study |
| pNTP2884 | a promoterless *lacZ* reporter plasmid | This study |
| pNTP2864:: PgtaT *-143 to +229* | Sequence from -143 to +229 of *C. crescentus* GTA cluster promoter PgtaT cloned into EcoRI-KpnI-cut pNTP2864:: PgtaT (-153 to +229) | This study |
| pNTP2864:: PgtaT-*133 to +229* | Sequence from -133 to +229 of *C. crescentus* GTA cluster promoter PgtaT cloned into EcoRI-KpnI-cut pNTP2864:: PgtaT (-153 to +229) | This study |
| pNTP2864:: PgtaT *-123 to +229* | Sequence from -123 to +229 of *C. crescentus* GTA promoter PgtaT cloned into EcoRI-KpnI-cut pNTP2864:: PgtaT (-153 to +229) | This study |
| pNTP2864:: PgtaT *-113 to +229* | Sequence from -113 to +229 of *C. crescentus* GTA cluster promoter PgtaT cloned into EcoRI-KpnI-cut pNTP2864:: PgtaT (-153 to +229) | This study |
| pNTP2864::PgtaT *-56 to +229* | Sequence from -56 to +229 of *C. crescentus* GTA cluster promoter PgtaT cloned into EcoRI-KpnI-cut pNTP2864:: PgtaT (-153 to +229) | This study |
| pNPTS138:: $ter_{GTA}$ | | This study |
| pNTP2884:: $ter_{GTA}$ | terGTA cloned into a promoterless lacZ reporter plasmid | This study |
| pNTP2864::PgtaT-*153 to+229* + $ter_{GTA}$ | terGTA was cloned between PgtaT-*153* to *+229* and lacZ | This study |
| pNPTS138::P*rsaA* | | This study |
| pNTP2864::P*rsaA* | | This study |
| pNTP2864::P*rsaA*+ $ter_{GTA}$ | terGTA was cloned between P*rsaA and lacZ* | This study |
| pNPTS138::$ter_{T1}$ | | This study |
| pNTP2884:: $ter_{T1}$ | *terT1* cloned into a promoterless *lacZ* reporter plasmid | This study |
| pNTP2864::PgtaT-*153 to+229* + $ter_{T1}$ | *terT1* was cloned between PgtaT-*153 to+229 and lacZ* | This study |
| pNTP2864::P*rsaA*+ $ter_{T1}$ | *terT1 was cloned between* P*rsaA* and *lacZ* | This study |
| pNPTS138::mut10-1 | PgtaT (-153 to +229) with a mutated -10 promoter sequence | This study |
| pNTP2864::mut10-1 | | This study |

| | | |
|---|---|---|
| pNPTS138::mut10-2 | PgtaT (-153 to +229) with a mutated -10 promoter sequence | This study |
| pNTP2864::mut10-2 | | This study |
| pNPTS138::mut35-1 | PgtaT (-153 to +229) with a mutated -35 promoter sequence | This study |
| pNTP2864::mut35-1 | | This study |
| pNPTS138::mut35-2 | PgtaT (-153 to +229) with a mutated -35 promoter sequence | This study |
| pNTP2864::mut35-2 | | This study |
| pNPTS138::Δ*gafZ* | To delete *gafZ* | This study |
| pNPTS138::Δ*rogA* | To delete *rogA* | This study |
| pMCS1-Tn5-ME-R6Kγ-kanamycin$^{R}$-ME | Tn5 delivery plasmid, spectinomycinR and kanamycinR, R6Kγ origin | [3] |

| Oligonucleotides | Sequence | Sources |
|---|---|---|
| **For the construction of pNPTS138::gtaM-lacA** | | |
| TLO3300 | GCAATTGAAGCCGGCTGGCGCCAGTCAGCATC GACGAGTGGCTGGCCGAAGAG | This study |
| TLO3301 | GGTTGGCCATGTTCGCTCCCTTCTTACGACGCC GCGAACTTCAGCAGCTTGATC | This study |
| TLO3302 | GAAGGGAGCGAACATGGCCAACCTGAACGGCC GCGCGCGTC | This study |
| TLO3303 | TCAGAGCCAGACCGCGAAGGGCGTCGCGTCGG TTCAGC | This study |
| TLO3304 | CGCCCTTCGCGGTCTGGCTCTGATCCAATCCTC CCCCTGACGGGGGAGGTGTCCG | This study |
| TLO3305 | CGGCCGAAGCTAGCGAATTCGTGCTCGATCCC CTCCCCGCCGACACCGCCGACC | This study |
| **For the construction of pNPTS138::flag-ihfA** | | |
| NTO2830 | GCAATTGAAGCCGGCTGGCGCCACGGCAAGCT CCGGATGCTGGGC | This study |
| NTO2831 | CTTGTCGTCATCGTCTTTGTAGTCCATGCAACCC CCTTCACGCGTATC | This study |
| NTO2832 | GACTACAAAGACGATGACGACAAGAAGGGCGC TACTTTGACCCG | This study |
| NTO2833 | CGGCCGAAGCTAGCGAATTCGTGCAGCAGGGT GCGGACGCCATTC | This study |
| **For the construction of pNPTS138::ihfB-flag** | | |

| | | |
|---|---|---|
| NTO2745 | GCAATTGAAGCCGGCTGGCGCCAGACTAAGCTT CGTCTTTCATAGACG | This study |
| NTO2746 | CTTGTCGTCATCGTCTTTGTAGTCTTCGTCGCC GTCGGCGTTCAGGC | This study |
| NTO2747 | GACTACAAAGACGATGACGACAAGTAGCAACAA TCATGACCCGGGCTTG | This study |
| NTO2748 | CGGCCGAAGCTAGCGAATTCGTGCCATGTTCGC TCCAGACCCGGATC | This study |
| **For the construction of pNPTS138::ΔihfA** | | |
| NTO2799 | GCAATTGAAGCCGGCTGGCGCCAGTTGCGGTC GCCGATGTCGACTG | This study |
| NTO2800 | GCCCTTCATGCAACCCCCTTCAC | This study |
| NTO2801 | GTGAAGGGGGTTGCATGAAGGGCGGCGGCTAA GGATCGATTGCGG | This study |
| NTO2802 | CGGCCGAAGCTAGCGAATTCGTGTGCCCTTGG GCAACACCGATTTG | This study |
| **For construction of pNPTS138::ΔihfB** | | |
| NTO2781 | GCAATTGAAGCCGGCTGGCGCCATCGAAGTTC GCTTCGGCGAAGAC | This study |
| NTO2782 | CTTGATCATCGATATCCCCGGAAC | This study |
| NTO2783 | GTTCCGGGGATATCGATGATCAAGGACGAATAG CAACAATCATGACC | This study |
| NTO2784 | CGGCCGAAGCTAGCGAATTCGTGTCCAGACCC GGATCAGCCCTTCTTG | This study |
| **For the construction of pXYFPC-1:: ihfA** | | |
| NTO2845 | CGCTCGAGTTTTGGGGAGACGACCATATGAAGG GCGCTACTTTGACCC | This study |
| NTO2846 | AGTGGATCCCCCGGGCTGCAGCTAGCTTAGCC GCCCAGAGCGCGATCGATG | This study |
| **For the construction of pXYFPC-1:: ihfB** | | |

| | | |
|---|---|---|
| NTO2767 | CGCTCGAGTTTTGGGGAGACGACCATATGATCA AGTCTGAACTCATCGCCAG | This study |
| NTO2768 | AGTGGATCCCCCGGGCTGCAGCTAGCCTATTC GTCGCCGTCGGCGTTCAG | This study |
| **For the construction of pNPTS138::ΔgtaT IBE** | | |
| NTO2230 | GCAATTGAAGCCGGCTGGCGCCA ACGCCGGCGAACAGGCGCAGC | This study |
| NTO2753 | CTTATTCAGAGCGTTCAGGCCCG | This study |
| NTO2754 | CGGGCCTGAACGCTCTGAATAAGATCATCCGCG CCCGCTGGCGCCG | This study |
| NTO2755 | CGGCCGAAGCTAGCGAATTCGTGAGCATCAGC CAGGTGGACCACGGGC | This study |
| **For the construction of pNPTS138::gtaT IBE (TT to GG)** | | |
| NTO2230 | GCAATTGAAGCCGGCTGGCGCCA ACGCCGGCGAACAGGCGCAGC | This study |
| NTO2863 | CCGCCGCTGATTTCGCGAGACTTC | This study |
| NTO2864 | GAAGTCTCGCGAAATCAGCGGCGGGTCACTTTT ACGCGCCCGCCAATC | This study |
| NTO2755 | CGGCCGAAGCTAGCGAATTCGTGAGCATCAGC CAGGTGGACCACGGGC | This study |
| **For the construction of pNPTS138::ΔgafY IBE** | | |
| NTO2749 | GCAATTGAAGCCGGCTGGCGCCACGGCCGCTC TATGGCCGTGAAGC | This study |
| NTO2750 | ATGAGGTTCTGTGGATAGAACGC | This study |
| NTO2751 | GCGTTCTATCCACAGAACCTCATCCCCGTGAGA CATGAGCGGAATAG | This study |
| NTO2752 | CGGCCGAAGCTAGCGAATTCGTGCCTCGAAATG GATCTGAAGCCC | This study |
| **For the construction of pNPTS138::gafY IBE (TT to GG)** | | |
| NTO2749 | GCAATTGAAGCCGGCTGGCGCCACGGCCGCTC TATGGCCGTGAAGC | This study |

| | | |
|---|---|---|
| NTO2865 | CTCCGCGCTTGTTTTTCCATCGCCTAGG | This study |
| NTO2866 | AGGCGATGGAAAAACAAGCGCGGAGCCCCGTG AGACATGAGCGGAATAGG | This study |
| NTO2752 | CGGCCGAAGCTAGCGAATTCGTGCCTCGAAATG GATCTGAAGCCC | This study |
| **For the construction of pNPTS138::Δihf500** | | |
| NTO2888 | GCAATTGAAGCCGGCTGGCGCCAGGCGGCGAC ATGGTCCGCTCGGTG | This study |
| NTO2889 | AGATTTGCCCCCGGCGAACACCTCTC | This study |
| NTO2890 | GAGGTGTTCGCCGGGGGCAAATCTCTCCCCCA GAGGAGGAGAGATGTTC | This study |
| NTO2891 | CGGCCGAAGCTAGCGAATTCGTGGGAAGGCGA AACTGAAACCCCCTC | This study |
| **For the construction of pNPTS138::ihfall4GG** | | |
| NTO2888 | GCAATTGAAGCCGGCTGGCGCCAGGCGGCGAC ATGGTCCGCTCGGTG | This study |
| NTO2892 | GTCGGCCTCTTCCCCGCCCTAACTGC | This study |
| NTO2893 | GCAGTTAGGGCGGGGAAGAGGCCGAC | This study |
| NTO2929 | CCCGCCCTAGCTGCGGGAGATTTTG | This study |
| NTO2932 | CAAAATCTCCCGCAGCTAGGGCGGGGAAGAGG CGCGTGATTTCTCCTC | This study |
| NTO2935 | CTCGGTGGATCGGCCGGCCACAG | This study |
| NTO2934 | CTGTGGCCGGCCGATCCACCGAGGGCTGAGAT CGCCG | This study |
| NTO2905 | TCACTCTCCTCTTCCACGCCCTAACTGCGGAAA ATTTGC | This study |
| NTO2899 | CGCAGTTAGGGCGTGGAAGAGGAGAGTGA | This study |
| NTO2891 | CGGCCGAAGCTAGCGAATTCGTGGGAAGGCGA AACTGAAACCCCCTC | This study |
| **For the construction of pNPTS138::nusG-vsvg** | | |

| | | |
|---|---|---|
| NTO2923 | GCAATTGAAGCCGGCTGGCGCCACAACCAGCT GCTCAAGCTGGCGACC | This study |
| NTO2924 | CTTACCCAGGCGGTTCATTTCGATATCAGTGTA GGCGATCTTTTCGACCTGATTG | This study |
| NTO2925 | TACACTGATATCGAAATGAACCGCCTGGGTAAG TGACGCACTTCAGGCCTGCTGAAATC | This study |
| NTO2926 | CGGCCGAAGCTAGCGAATTCGTGCACCACCCA GCCCATGGCCGGGCTG | This study |
| **For the construction of pNPTS138::vsvg-nusA** | | |
| NTO4041 | GCAATTGAAGCCGGCTGGCGCCATTCGCCTGC GCCTGATGGGCGG | This study |
| NTO4042 | CTTACCCAGGCGGTTCATTTCGATATCAGTGTA CATGGTCAGTCCTCTTCACTTTC | This study |
| NTO4043 | ATCGAAATGAACCGCCTGGGTAAGGCCATCGG CATCGCCGCCAACCG | This study |
| NTO4044 | CGGCCGAAGCTAGCGAATTCGTGGCGCGGATG CGATCGCCGACATTG | This study |
| **For the construction of pNPTS138::vsvg-nusE** | | |
| NTO4037 | GCAATTGAAGCCGGCTGGCGCCACCCGTGAGG AAGCACGTCGGCGG | This study |
| NTO4038 | CTTACCCAGGCGGTTCATTTCGATATCAGTGTA CATATCGCTCGGTTTCCTAGTGG | This study |
| NTO4039 | ATCGAAATGAACCGCCTGGGTAAGGATCGTCAG AACATCCGCATCCGGC | This study |
| NTO4040 | CGGCCGAAGCTAGCGAATTCGTGGGAACGTAC GACCCGGGTCCTGG | This study |
| **For the construction of pNPTS138::flag-rpoD** | | |
| NTO4268 | GCAATTGAAGCCGGCTGGCGCCATGGAGTGAA AGCGGCGGTCCCGAGG | This study |
| NTO4269 | cttgtcgtcatcgtctttgtagtcCATCAAGCTCTCCGAAGTCGACGC G | This study |

| | | |
|---|---|---|
| NTO4270 | gactacaaagacgatgacgacaagAGCAACAATTCCTCGGCCGAGAC | This study |
| NTO4271 | CGGCCGAAGCTAGCGAATTCGTGGATCGCGGCGTAGGTGCCTTCC | This study |
| **For the construction of pCOLADuet-1::6xhis gafZ** | | |
| NTO2693 | CCATCATCACCACAGCCAGGATCCGAATTCGATGACCGGCGGGCTTCAGATCC | This study |
| NTO2694 | TCGACTTAAGCATTATGCGGCCGCAAGCTTTCATGCCATCCGGTAGTGTCGG | This study |
| **For the construction of pCOLADuet-1:: 6xhis gafZ gafY** | | |
| NTO2691 | AGTTAAGTATAAGAAGGAGATATACATATGCTGGTTCAAGCCCTTGAGGAC | This study |
| NTO2692 | AGCGGTTTCTTTACCAGACTCGAGGGTACCTTACGGCAGATCCTGAGCGTGCG | This study |
| **For the construction of pCOLADuet-1::6xhis rpoD gafY** | | |
| NTO4289 | CCATCATCACCACAGCCAGGATCCGAATTCGATGAGCAACAATTCCTCGGCCG | This study |
| NTO4290 | TCGACTTAAGCATTATGCGGCCGCAAGCTTTTACGAGTCCAGGAAGCTGCGC | This study |
| **For the construction of pCOLADuet-1::6xhis rpoD (domain 3+4)** | | |
| NTO4349 | CCATCATCACCACAGCCAGGATCCGAATTCGATGATCCGCATCCCGGTGCACATGATC | This study |
| NTO4290 | TCGACTTAAGCATTATGCGGCCGCAAGCTTTTACGAGTCCAGGAAGCTGCGC | This study |
| **For the construction of pNTP2846** | | |
| NTO4188 | ACTAGTTCTAGAGCGGCCGTCGACGG | This study |
| NTO4189 | CCGTCGACGGCCGCTCTAGAACTAGTCAAATAAAACGAAAGGCTCAGTCG | This study |
| NTO4190 | CCCGGGCTGCAGCTAGCTTACTTG | This study |

| | | |
|---|---|---|
| NTO4191 | CAAGTAAGCTAGCTGCAGCCCGGGACTAGTTCTAGAGCGGCCATTCAC | This study |
| **For the construction of pNTP2864** | | |
| NTO4192 | ATATAAAAACTGTTGTAATTCATTAAGC | This study |
| NTO4193 | GCTTAATGAATTACAACAGTTTTTATATAGCGCTACCGGACTCAGATCCAGATC | This study |
| NTO4194 | GCTAGCTTACTTGTACAGCTCGTCCATG | This study |
| NTO4195 | CATGGACGAGCTGTACAAGTAAGCTAGC | This study |
| **For the construction of pRlacZ290:: PgtaT (-153 to +229)** | | |
| NTO4085 | CGGATCGCACGAACCCGCTGAATGGGAATTCTTGCGGCGCGCTTGCCCGTCAG | This study |
| NTO4101 | GCAGGTCGACTCTAGAGGATCCCCGGGTACCCGGCCGCCTCGTCGGCGGCGGGTTTG | This study |
| **For the construction of pNTP2864:: PgtaT (-153 to +229)** | | |
| NTO4196 | CGCTCGAGTTTTGGGGAGACGACCATATGGAATTCTTGCGGCGCGCTTGCCCGTCAG | This study |
| NTO4197 | AGTGGATCCCCCGGGCTGCAGCTAGCTTATTTTTGACACCAGACCAACTGG | This study |
| **For the construction of pNPTS138:: PgtaT (-113 to +229)** | | |
| NTO4259 | GCAATTGAAGCCGGCTGGCGCCAGAATTCTTGGCGGGCCTGAACGCTCTGAATAAG | This study |
| NTO4185 | CGGCCGAAGCTAGCGAATTCGTGGGTACCCGGCCGCCTCGTCGGCGGCGGGTTTG | This study |
| **For the construction of pNPTS138:: PgtaT (-56 to +229)** | | |
| NTO4369 | GCAATTGAAGCCGGCTGGCGCCAGAATTCGTCACTTTTACGCGCCCGCCAATC | This study |

| | | |
|---|---|---|
| NTO4185 | CGGCCGAAGCTAGCGAATTCGTGGGTACCCGG CCGCCTCGTCGGCGGCGGGTTTG | This study |
| **For the construction of pNPTS138:: PgtaT (-143 to +229)** | | |
| NTO4256 | GCAATTGAAGCCGGCTGGCGCCAGAATTCCTTG CCCGTCAGGGGCCTCGAC | This study |
| NTO4185 | CGGCCGAAGCTAGCGAATTCGTGGGTACCCGG CCGCCTCGTCGGCGGCGGGTTTG | This study |
| **For the construction of pNPTS138:: PgtaT (-133 to +229)** | | |
| NTO4257 | GCAATTGAAGCCGGCTGGCGCCAGAATTCAGG GGCCTCGACTATGGGGATTG | This study |
| NTO4185 | CGGCCGAAGCTAGCGAATTCGTGGGTACCCGG CCGCCTCGTCGGCGGCGGGTTTG | This study |
| **For the construction of pNPTS138:: PgtaT (-123 to +229)** | | |
| NTO4258 | GCAATTGAAGCCGGCTGGCGCCAGAATTCACTA TGGGGATTGGCGGGCCTGAAC | This study |
| NTO4185 | CGGCCGAAGCTAGCGAATTCGTGGGTACCCGG CCGCCTCGTCGGCGGCGGGTTTG | This study |
| **For the construction of pNTPS138:: Δ125bp (+90 to +214)** | | |
| NTO2230 | GCAATTGAAGCCGGCTGGCGCCA ACGCCGGCGAACAGGCGCAGC | This study |
| NTO2764 | AAGGCGGCGATGAGCTTGTCCATC | This study |
| NTO2765 | GATGGACAAGCTCATCGCCGCCTTCCGACGAG GCGGCCGATGATGAC | This study |
| NTO2766 | CGGCCGAAGCTAGCGAATTCGTGTTCGGCCAG GCGCAGAACTCGTC | This study |

| | | |
|---|---|---|
| **For the construction of pNTPS138:: Δ144bp (+230 to 373)** | | |
| NTO2957 | GCAATTGAAGCCGGCTGGCGCCAGCCCGTCAG ATGCGCCATGGTC | This study |
| NTO2958 | CGGCCGCCTCGTCGGCGGCGGGTTTG | This study |
| NTO2960 | AAACCCGCCGCCGACGAGGCGGCCGAGCCTG GAGCCGGAGGATCAAC | This study |
| NTO2961 | CGGCCGAAGCTAGCGAATTCGTGGAAGGCGGG CGCCAGGTTGCCGG | This study |
| **For the construction of pNPTS138::gtaT (ATG1 to TGA)** | | |
| NTO2957 | GCAATTGAAGCCGGCTGGCGCCAGCCCGTCAG ATGCGCCATGGTC | This study |
| NTO2962 | TCACGGCCGCCTCGTCGGCGGC | This study |
| NTO2963 | GCCGCCGACGAGGCGGCCGTGAATGACGAGG CTTCCATGCATGACG | This study |
| NTO2961 | CGGCCGAAGCTAGCGAATTCGTGGAAGGCGGG CGCCAGGTTGCCGG | This study |
| **For the construction of pNPT138::gtaT (ATG2 to TGA)** | | |
| NTO2957 | GCAATTGAAGCCGGCTGGCGCCAGCCCGTCAG ATGCGCCATGGTC | This study |
| NTO2964 | TCACATCGGCCGCCTCGTCGGC | This study |
| NTO2965 | GCCGACGAGGCGGCCGATGTGAACGAGGCTTC CATGCATGACG | This study |
| NTO2961 | CGGCCGAAGCTAGCGAATTCGTGGAAGGCGGG CGCCAGGTTGCCGG | This study |
| **For the construction of pNPTS138::gtaT (ATG3 to TGA)** | | |
| NTO2957 | GCAATTGAAGCCGGCTGGCGCCAGCCCGTCAG ATGCGCCATGGTC | This study |
| NTO2966 | TCAGCATGGAAGCCTCGTCATC | This study |
| NTO2967 | GATGACGAGGCTTCCATGCTGAACGGCATTCCG GACGACCCCG | This study |

| | | |
|---|---|---|
| NTO2961 | CGGCCGAAGCTAGCGAATTCGTGGAAGGCGGG CGCCAGGTTGCCGG | This study |
| **For the construction of pNPTS138:: Δ336bp (-113 to +223)** | | |
| NTO2230 | GCAATTGAAGCCGGCTGGCGCCA ACGCCGGCGAACAGGCGCAGC | This study |
| NTO2906 | TCCCCATAGTCGAGGCCCCTGACG | This study |
| NTO2907 | CGTCAGGGGCCTCGACTATGGGGACGGCCGAT GATGACGAGGCTTCC | This study |
| NTO2766 | CGGCCGAAGCTAGCGAATTCGTGTTCGGCCAG GCGCAGAACTCGTC | This study |
| **For the construction of pNPTS138::gtaT (TTG to CTG)** | | |
| NTO2230 | GCAATTGAAGCCGGCTGGCGCCA ACGCCGGCGAACAGGCGCAGC | This study |
| NTO2908 | GCGTTCAGGCCCGCCAGTCCCCATAGTCGAGG | This study |
| NTO2909 | CCTCGACTATGGGGACTGGCGGGCCTGAACGC | This study |
| NTO2766 | CGGCCGAAGCTAGCGAATTCGTGTTCGGCCAG GCGCAGAACTCGTC | This study |
| **For the construction of pNPTS138::Δ*YBE*** | | |
| NTO2230 | GCAATTGAAGCCGGCTGGCGCCA ACGCCGGCGAACAGGCGCAGC | This study |
| NTO2970 | GGGCAAGCGCGCCGCAAAAACCG | This study |
| NTO2971 | CGGTTTTTGCGGCGCGCTTGCCCGGGATTGGC GGGCCTGAACGCTC | This study |
| NTO2755 | CGGCCGAAGCTAGCGAATTCGTGAGCATCAGC CAGGTGGACCACGGGC | This study |
| **pNPTS138:: YBE (ACTATG to TGACCC)** | | |
| NTO2230 | GCAATTGAAGCCGGCTGGCGCCA ACGCCGGCGAACAGGCGCAGC | This study |
| NTO4157 | GGGTCACGAGGCCCCTGACGGGCAAGCGCGC CGCAAAAACCG | This study |
| NTO4158 | GTCAGGGGCCTCGTGACCCGGGATTGGCGGGC CTGAACGCTC | This study |
| NTO2755 | CGGCCGAAGCTAGCGAATTCGTGAGCATCAGC CAGGTGGACCACGGGC | This study |
| **For the construction of pNPTS138::ΔGTA core promoter** | | |
| NTO2230 | GCAATTGAAGCCGGCTGGCGCCA ACGCCGGCGAACAGGCGCAGC | This study |

| | | |
|---|---|---|
| NTO2760 | GGCGCGGATGATTGGCGGGCGCGTAAAAG | This study |
| NTO2761 | ACGCGCCCGCCAATCATCCGCGCCACGAGGTC CGGGACTGGAGCGC | This study |
| NTO2755 | CGGCCGAAGCTAGCGAATTCGTGAGCATCAGC CAGGTGGACCACGGGC | This study |
| **For the construction of pNPTS138:: ZBE* (CTGGCGC to TTTTTTC)** | | This study |
| NTO2230 | GCAATTGAAGCCGGCTGGCGCCA ACGCCGGCGAACAGGCGCAGC | This study |
| NTO4090 | CGCTTCTTCCGGCCATGACACAAAGATACGGAA AAAACGGGCGCGGATGATTGGCGGGCGCG | This study |
| NTO4091 | CGTTTTTTCCGTATCTTTGTGTCATGGCCGGAAG AAGCGACGAGGTCCGGGACTGGAGCGCGACGC | This study |
| NTO2755 | CGGCCGAAGCTAGCGAATTCGTGAGCATCAGC CAGGTGGACCACGGGC | This study |
| **For the construction of pNPTS138::mut10-1** | | |
| NTO4184 | GCAATTGAAGCCGGCTGGCGCCAGAATTCTTGC GGCGCGCTTGCCCGTCAGG | This study |
| NTO4174 | GGGGGGGCGGCGCCAGCGGGCGCGGATGATT G | This study |
| NTO4175 | CAATCATCCGCGCCCGCTGGCGCCGCCCCCCC GTGTCATGGCCGGAAGAAGCGAC | This study |
| NTO4185 | CGGCCGAAGCTAGCGAATTCGTGGGTACCCGG CCGCCTCGTCGGCGGCGGGTTTG | This study |
| **For the construction of pNPTS138::mut10-2** | | |
| NTO4184 | GCAATTGAAGCCGGCTGGCGCCAGAATTCTTGC GGCGCGCTTGCCCGTCAGG | This study |
| NTO4176 | GGGGAGATACGGCGCCAGCGGGCGCGGATG | This study |
| NTO4177 | CATCCGCGCCCGCTGGCGCCGTATCTCCCCGT CATGGCCGGAAGAAGCGACGAGGTC | This study |
| NTO4185 | CGGCCGAAGCTAGCGAATTCGTGGGTACCCGG CCGCCTCGTCGGCGGCGGGTTTG | This study |
| **For the construction of pNPTS138::mut35-1** | | |

| | | |
|---|---|---|
| NTO4184 | GCAATTGAAGCCGGCTGGCGCCAGAATTCTTGC GGCGCGCTTGCCCGTCAGG | This study |
| NTO4242 | GGGGGGGGCGGGCGCGTAAAAGTGAC | This study |
| NTO4243 | GTCACTTTTACGCGCCCGCCCCCCCCCGCGC CCGCTGGCGCCGTATCTTTG | This study |
| NTO4185 | CGGCCGAAGCTAGCGAATTCGTGGGTACCCGG CCGCCTCGTCGGCGGCGGGTTTG | This study |
| **For the construction of pNPTS138::mut35-2** | | |
| NTO4184 | GCAATTGAAGCCGGCTGGCGCCAGAATTCTTGC GGCGCGCTTGCCCGTCAGG | This study |
| NTO4264 | CCCTTGGCGGGCGCGTAAAAGTG | This study |
| NTO4265 | CACTTTTACGCGCCCGCCAAGGGTCCGCGCCC GCTGGCGCCGTATC | This study |
| NTO4185 | CGGCCGAAGCTAGCGAATTCGTGGGTACCCGG CCGCCTCGTCGGCGGCGGGTTTG | This study |
| **For the construction of pNPTS138::P*rsaA*** | | |
| NTO4252 | GCAATTGAAGCCGGCTGGCGCCAGAATTCTGCT GTACCGGTTAGAAAAATGCTG | This study |
| NTO4253 | CGGCCGAAGCTAGCGAATTCGTGGGTACCGCA TGGGAGCAGCGATGGCTATAG | This study |
| **For the construction of pNPTS138::$ter_{GTA}$** | | |
| NTO4094 | GCAATTGAAGCCGGCTGGCGCCAGGATCCGGG CGGCCGAAGTAACTCTCCTC | This study |
| NTO2930 | CTCTTCAACGCCCTAACTGCGGG | This study |
| NTO2931 | CCCGCAGTTAGGGCGTTGAAGAG | This study |
| NTO2935 | CTCGGTGGATCGGCCGGCCACAG | This study |
| NTO2936 | CTGTGGCCGGCCGATCCACCGAG | This study |
| NTO4095 | CGGCCGAAGCTAGCGAATTCGTGAAGCTTGGG AAATCACTCTCCTCTTCAAC | This study |
| **For the construction of pNPTS138::$ter_{T1}$** | | |

| | | |
|---|---|---|
| NTO4151 | GCAATTGAAGCCGGCTGGCGCCAGGATCCATAAAACGAAAGGCTCAGTCGAAAG | This study |
| NTO4152 | CGGCCGAAGCTAGCGAATTCGTGAAGCTTAAACCACGACCTGGACCTCTTGCC | This study |
| **For the construction of pNPTS138:: Δ*gafZ*** | | |
| NTO2634 | GCAATTGAAGCCGGCTGGCGCCAAGGGGGCGATATCCTGCGTTCTATCC | This study |
| NTO2635 | GCCGGTCATTACGGCAGATCCTG | This study |
| NTO2636 | CAGGATCTGCCGTAATGACCGGCATGGCATGAGGCGACGTTCAGTC | This study |
| NTO2637 | CGGCCGAAGCTAGCGAATTCGTGAGGTGGCGGTCGGGCGCTTCGAG | This study |
| **For the construction of pNPTS138:: Δ*rogA*** | | |
| NTO2194 | GCAATTGAAGCCGGCTGGCGCCAGATGTCGTCGCGCTGTTGCTCC | This study |
| NTO2195 | CCAGATTTCGCCGTGAGACAG | This study |
| NTO2196 | CCCTGTCTCACGGCGAAATCTGGAGCCAGTAGGCCGACCCGACAG | This study |
| NTO2197 | CGGCCGAAGCTAGCGAATTCGTGGCGCTGATCGAGGAAAAGCTG | This study |
| **For the construction of pET15b::*gafY*** | | |
| NTO4346 | CTTTAAGAAGGAGATATACCATGGGCATGCTGGTTCAAGCCCTTGAGGACATC | This study |
| NTO4347 | TTCGGGCTTTGTTAGCAGCCGGATCCTTACGGCAGATCCTGAGCGTGCGAC | This study |

**Supplementary Table 3.** List of ChIP-seq data used in this study

| **ChIP-seq samples** | **Genetic backgrounds** | **antibody** | **replicates** | **Source** |
|---|---|---|---|---|
| sample 1 | CB15N WT | anti-FLAG | replicate 1 | This study |
| sample 2 | CB15N WT | anti-FLAG | replicate 2 | This study |

| | | | | |
|---|---|---|---|---|
| sample 3 | *CB15N ihfB::ihfB-flag* | anti-FLAG | replicate 1 | This study |
| sample 4 | *CB15N ihfB::ihfB-flag* | anti-FLAG | replicate 2 | This study |
| sample 5 | *∆rogA* | anti-FLAG | replicate 1 | This study |
| sample 6 | *∆rogA* | anti-FLAG | replicate 2 | This study |
| sample 7 | *∆rogA ihfB::ihfB-flag* | anti-FLAG | replicate 1 | This study |
| sample 8 | *∆rogA ihfB::ihfB-flag* | anti-FLAG | replicate 2 | This study |
| sample 9 | *ihfA::ihfA-flag* | anti-FLAG | replicate 1 | This study |
| sample 10 | *ihfA::ihfA-flag* | anti-FLAG | replicate 2 | This study |
| sample 11 | *∆rogA ihfA::ihfA-flag* | anti-FLAG | replicate 1 | This study |
| sample 12 | *∆rogA ihfA::ihfA-flag* | anti-FLAG | replicate 2 | This study |
| sample 13 | *∆rogA gtaT IBE (TT to GG)* | anti-FLAG | replicate 1 | This study |
| sample 14 | *∆rogA gtaT IBE (TT to GG)* | anti-FLAG | replicate 2 | This study |
| sample 15 | *∆rogA gafY IBE (TT to GG)* | anti-FLAG | replicate 1 | This study |
| sample 16 | *∆rogA gafY IBE (TT to GG)* | anti-FLAG | replicate 2 | This study |
| sample 17 | *∆rogA gtaT IBE (TT to GG) ihfB::ihfB-flag* | anti-FLAG | replicate 1 | This study |
| sample 18 | *∆rogA gtaT IBE (TT to GG) ihfB::ihfB-flag* | anti-FLAG | replicate 2 | This study |
| sample 19 | *∆rogA gafY IBE (TT to GG) ihfB::ihfB-flag* | anti-FLAG | replicate 1 | This study |
| sample 20 | *∆rogA gafY IBE (TT to GG) ihfB::ihfB-flag* | anti-FLAG | replicate 2 | This study |
| sample 21 | *∆rogA gtaT IBE2-3-4-5 (TT to GG)* | anti-FLAG | replicate 1 | This study |
| sample 22 | *∆rogA gtaT IBE2-3-4-5 (TT to GG)* | anti-FLAG | replicate 2 | This study |
| sample 23 | *∆rogA gtaT IBE2-3-4-5 (TT to GG) ihfB::ihfB-flag* | anti-FLAG | replicate 1 | This study |
| sample 24 | *∆rogA gtaT IBE2-3-4-5 (TT to GG) ihfB::ihfB-flag* | anti-FLAG | replicate 2 | This study |
| sample 25 | *∆rogA* | anti-GafY | replicate 1 | Gozzi *et al* (2022)[4] |
| sample 26 | *∆rogA* | anti-GafY | replicate 2 | Gozzi *et al* (2022)[4] |
| sample 27 | *∆rogA ∆gafY* | anti-GafY | replicate 1 | Gozzi *et al* (2022)[4] |

| | | | | |
|---|---|---|---|---|
| sample 28 | Δ*rogA* Δ*gafY* | anti-GafY | replicate 2 | Gozzi *et al* (2022)[4] |
| sample 29 | Δ*rogA* | anti-FLAG | replicate 1 | Gozzi *et al* (2022)[4] |
| sample 30 | Δ*rogA* | anti-FLAG | replicate 2 | Gozzi *et al* (2022)[4] |
| sample 31 | Δ*rogA gafZ::gafZ-FLAG* | anti-FLAG | replicate 1 | Gozzi *et al* (2022)[4] |
| sample 32 | Δ*rogA gafZ::gafZ-FLAG* | anti-FLAG | replicate 2 | Gozzi *et al* (2022)[4] |
| sample 33 | Δ*rogA rpoC::rpoC-3xflag* | anti-FLAG | replicate 1 | This study |
| sample 34 | Δ*rogA rpoC::rpoC-3xflag* | anti-FLAG | replicate 2 | This study |
| sample 35 | Δ*rogA gafZ::gafZ-FLAG nusA::vsvg-nusA* | anti-VSVG | replicate 1 | This study |
| sample 36 | Δ*rogA gafZ::gafZ-FLAG nusG::nusG-vsvg* | anti-VSVG | replicate 1 | This study |
| sample 37 | Δ*rogA gafZ::gafZ-FLAG nusE::vsvg-nusE* | anti-VSVG | replicate 1 | This study |
| sample 38 | Δ*rogA gafZ::gafZ-FLAG* | anti-VSVG | replicate 1 | This study |
| sample 39 | Δ*rogA gafZ::gafZ-FLAG nusA::vsvg-nusA* | anti-VSVG | replicate 2 | This study |
| sample 40 | Δ*rogA gafZ::gafZ-FLAG nusG::nusG-vsvg* | anti-VSVG | replicate 2 | This study |
| sample 41 | Δ*rogA gafZ::gafZ-FLAG nusE::vsvg-nusE* | anti-VSVG | replicate 2 | This study |
| sample 42 | Δ*rogA gafZ::gafZ-FLAG* | anti-VSVG | replicate 2 | This study |
| sample 43 | Δ*rogA gafZ::gafZ-FLAG ZBE** | anti-FLAG | replicate 1 | This study |
| sample 44 | Δ*rogA gafZ::gafZ-FLAG ZBE** | anti-FLAG | replicate 2 | This study |
| sample 45 | Δ*rogA gafZ::gafZ-FLAG ZBE** | anti-GafY | replicate 1 | This study |
| sample 46 | Δ*rogA gafZ::gafZ-FLAG ZBE** | anti-GafY | replicate 2 | This study |
| sample 47 | Δ*rogA YBE** | anti-GafY | replicate 1 | This study |
| sample 48 | Δ*rogA YBE** | anti-GafY | replicate 2 | This study |
| sample 49 | Δ*rogA rpoD::flag-rpoD* | anti-FLAG | replicate 1 | This study |
| sample 50 | Δ*rogA rpoD::flag-rpoD* | anti-FLAG | replicate 2 | This study |
